# COAL_PHYRE: A Composite Likelihood Method for Estimating Species Tree Parameters from Genomic Data Using Coalescent Theory

**DOI:** 10.1101/2020.11.17.387399

**Authors:** Geno Guerra, Rasmus Nielsen

**Affiliations:** Department of Statistics, University of California, Berkeley, California 94720, USA; Department of Integrative Biology, University of California, Berkeley, California 94720 USA; Lundbeck Foundations Centre for GeoGenetics, University of Copenhagen, Denmark

**Keywords:** phylogenetics, species tree, maximum likelihood, composite likelihood, mutational variance, gene tree, estimation error, multispecies coalescent, incomplete lineage sorting, genomic data, sequence data

## Abstract

Genome-scale data are increasingly being used to infer phylogenetic trees. A major challenge in such inferences is that different regions of the genome may have local topologies that differ from the species tree due to incomplete lineage sorting (ILS). Another source of gene tree discrepancies is estimation errors arising from the randomness of the mutational process during sequence evolution. There are two major groups of methods for estimating species tree from whole-genome data: a set of full likelihood methods, which model both sources of variance, but do not scale to large numbers of independent loci, and a class of faster approximation methods which do not model the mutational variance.

To bridge the gap between these two classes of methods, we present COAL_PHYRE (COmposite Approximate Likelihood for PHYlogenetic REconstruction), a composite likelihood based method for inferring population size and divergence time estimates of rooted species trees from aligned gene sequences. COAL_PHYRE jointly models coalescent variation across loci using the MSC and variation in local gene tree reconstruction using a normal approximation. To evaluate the accuracy and speed of the method, we compare against BPP, a powerful MCMC full-likelihood method, as well as ASTRAL-III, a fast approximate method. We show that COAL_PHYRE’s divergence time and population size estimates are more accurate than ASTRAL, and comparable to those obtained using BPP, with an order of magnitude decrease in computational time. We also present results on previously published data from a set of Gibbon species to evaluate the accuracy in topology and parameter inference on real data, and to illustrate the method’s ability to analyze data sets which are prohibitively large for MCMC methods.

## 3 Introduction

With the continued improvement of sequencing technologies, inferring evolutionary relationships between organisms using multi-gene sequences has become the standard in the field of phylogenetics. Bifurcating species trees are a common way to represent these relationships, with branching points representing speciation events. While a species tree represents the history of these species as a whole, trees in individual genome segments can have their own, potentially discordant, topology due to horizontal gene transfer, gene duplication/loss, and/or incomplete lineage sorting (ILS) [21]. The most ubiquitous of these, ILS, is of particular focus in the field [5], and can be well-modeled using the multi-species coalescent (MSC) (see e.g., [27]). Many methods exist to infer the species tree topology of a group of organisms using the MSC in the presence of ILS, and are shown to be statistically consistent assuming the gene tree topologies are known without error [15, 23, 20]. This assumption however is unrealistic, as gene trees typically are estimated from sequence data, with a finite amount of mutations present. The random process of mutation adds a second layer of variation among gene trees, and ignoring this can lead to poor method performance [10, 11, 14]. A class of Bayesian hierarchical methods exist, which jointly model gene and species tree topologies in a full likelihood framework (e.g. [17, 4, 19, 6, 9, 34), and account for both coalescent and mutational variance, but these approaches have been shown to be computationally intensive (100s of hours) and not able to scale to large amounts of genes or species [18, 22, 31].

Although it has been known for decades that gene trees can differ in topology from an underlying species tree, a common approach to estimating trees and divergence times to avoid gene tree estimation error still relies on concatenated “super-matrices” of gene sequences (where multiple gene alignments are concatenated together to form one large “super gene”). Under high levels of mutational variation, this concatenation approach was justified as a way to pool information between highly noisy genes. [32, 16, 8] discuss results showing that concatenation-based approaches are not always outperformed by more ILS-sensitive methods. In short, concatenation methods seem to be predictably less accurate than coalescent based methods under high ILS (when there are short branches in the true species tree) and can even give high confidence to incorrect topologies [29]. Away from these scenarios, concatenation can empirically perform equal to or better than coalescent based methods. As such, concatenation is still widely used for inferring phylogenies in many empirical studies.

Divergence time estimates have become an essential addition in phylogenetic inference, as many studies utilize or require time-calibrated phylogenies, for example in biogeography, or in modeling of character evolution [3, 28, 25]. In particular, a challenging problem in phylogenetics is accurately inferring divergence times and population sizes in the presence of mutational variance. The Bayesian method, BPP [34] provides highly accurate results under the assumption of a molecular clock and the Jukes and Cantor model of sequence evolution [13]. However, this method, along with other Bayesian approaches, is unable to take advantage of the full information in genomic data sets, and must instead subdivide data into smaller (~ 100 segments) blocks of genes per run to perform inference in reasonable amounts of time.

In this paper, we present a coalescent based method to jointly infer species divergence times and ancient population sizes in the presence of mutational variance/gene tree estimation error. For a given topology, or set of *k* topologies, our method COAL-PHYRE (COmposite Approximate Likelihood for PHYlogenetic REconstruction) uses a composite likelihood approach to estimate tree parameters from DNA sequence data. COAL-PHYRE is able to analyze data with tens of thousands of genes/loci and multiple individuals in each sampled species. We show that the divergence time and population size estimates of COAL-PHYRE are comparable to the more time intensive estimates obtained using BPP [34], with at least an order of magnitude decrease in run time. We also compare to the popular approximate likelihood method ASTRAL-III [35], to compare the accuracy of our method against one that does not directly model mutational variance. Lastly, we analyze a data set of Gibbon species previously analyzed by BPP in [30], and find highly similar estimated parameters.

## 4 Methods

We consider a rooted bifurcating species tree 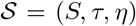 parameterized by topology *S*, divergence times *τ*, and population sizes *η*. See figure 1(a) for an illustration. Given a recombination-free region of the genome, *l* (interchangeably referred to as a gene or locus throughout), it is expected that that species tree topology *S* and the true local gene tree 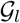 will not always match due to incomplete lineage sorting (ILS), which is common when branch lengths are short relative to the effective population sizes. Let 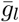 represent an estimated rooted topology with branch lengths of the local ancestry from the region l. Note that 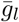 need not be bifurcating if the available genetic data is unable to resolve splits in the tree. This reconstructed gene tree is an estimate of the true local relationship between individuals, 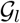. For any finite amount of information, (number of pairwise mutational differences on l), there is estimation variance in 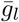. If 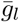 was known without error, meaning 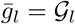, the MSC can be used to completely model the variation within and across gene trees, such as in STEM [15]. In reality, however, 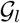 cannot be reliably estimated without sampling variance, and accurate estimation of the species tree from a collection of estimated gene trees requires models accounting for both the distribution of 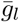 given 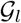, and 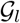 given 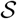.

**Figure 1:**
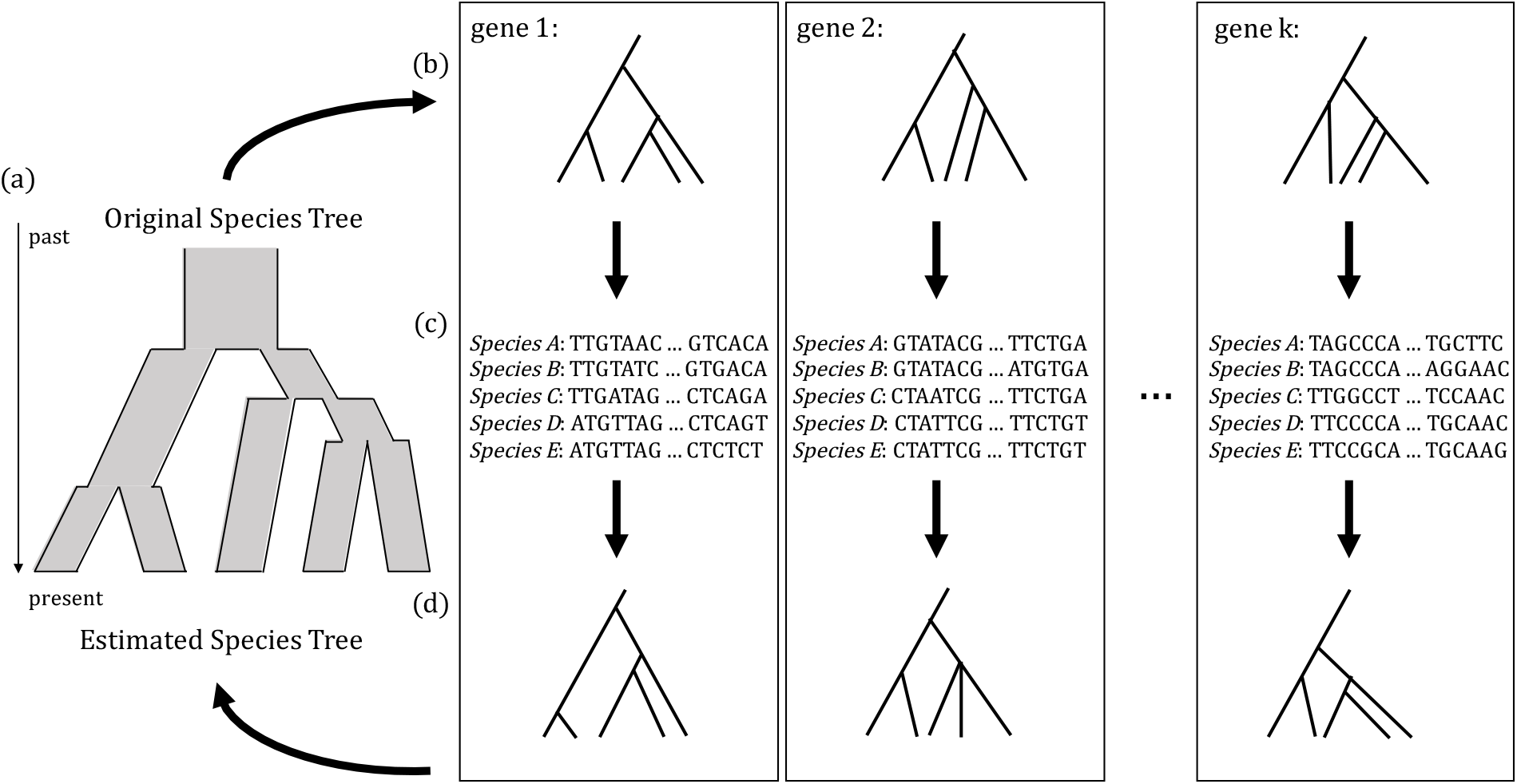
Contribution of coalescent and mutational variance. (a) Original bifurcating species tree. (b) *K* gene trees, each a different realization of a stochastic lineage sorting process on the original species tree. (c) Sequences created from the mutational process on each gene tree. (d) Gene trees estimated from the sequence data, which can differ in topology and branch length from the true gene trees due to mutational variance.

Our goal is to incorporate the effect of mutational variance directly into the likelihood in an interpretable way that is computationally tractable and scalable to many genes. We propose studying the observed distribution of individual coalescence times to do this.

We use the approximation that ‘noisy’ coalescence times (coalescence times estimated with mutational variation present) are well approximated by a hierarchical model of the MSC with an added normal distribution to capture both the coalescent and mutational variance, respectively. When coalescence times are estimated from sequence data, the layer of noise from gene tree reconstruction error (mutational variance) effectively smooths out the exponential-like distribution of the MSC, and the times fit closely to this hierarchical model.

Our method takes as input a set of aligned sequence data, and a rooted species tree topology (or set of topologies), and returns the inferred divergence times and population sizes which maximize the composite likelihood of pairwise coalescence times across the inputted loci, along with a likelihood, for each inputted topology. We assume there is no recombination within a locus, and allow free recombination between loci, and therefore assume loci are independent. To make use of the MSC, we assume the sequences have evolved on the gene tree under a molecular clock. Although not the goal of this paper, mutation rate variation between species can be incorporated into the gene tree estimation process if the computed gene trees have time measured in some real-time units as this satisfies the ultrametric property. We model each estimated pairwise coalescence time at a locus as an independent draw from a hierarchical MSC-normal distribution. The distribution of true coalescence times is modeled by the MSC, under a proposed species tree 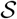. Conditional on those times, the normal distribution is then parameterized using the approximated mutational variance, derived from properties of the Poisson distribution. Our goal is to infer a set of divergence times and population sizes that maximize the composite likelihood of the estimated gene trees.

### 4.1 Mutational variance

As is common in most species tree inference methods ([34, 23, 15] for example), we assume that genomic data can be divided into recombination-free regions, with free recombination between regions. At any given locus, l, the underlying true gene tree 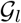 (including branch lengths) is not known but can be estimated from aligned sequence data. This estimated gene tree 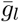 is a topology with estimated coalescence times.

For a specific time on the estimated tree, we can decompose the estimated time 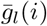 into a mixture of two components: the true coalescent time 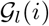, and then the estimation error resulting from having only a finite number mutations on each branch e_l_(i) (see figure 1). Mathematically, we can write this as:

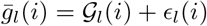

We approximate that error *ϵ_l_*(*i*), the difference between the estimated and the (unknown) true coalescence time, as distributed with mean 0 and variance *ξ_l_*(*i*), i.e., we assume that an unbiased estimator has been used to estimate 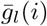. While 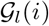 can be modeled using the MSC, we use the Poisson distribution of mutations given a coalescence time to quantify the variance *ξ_l_*(*i*), meaning *ξ_l_*(*i*) is a function of the unknown true coalescence time 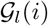.

Under the infinite sites assumption, the number of mutations on a lineage is Poisson distributed and the variance in the estimate of the coalescence time will also follow that of a Poisson distribution. In real life applications, the divergence between sequences is often estimated using finite-sites models. However, even for these models the Poisson variance might be a reasonable approximation, and we will evaluate the performance of all estimators presented in this paper using simulations under finite sites models. The estimation variance from the mutation process is then

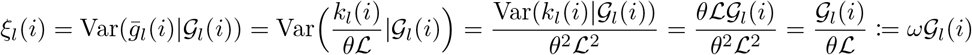

where 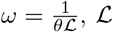 is the length of locus l, and *k_l_*(*i*) is the number of pairwise mutations for i on locus *l*.

While using the variance from the Poisson, we will approximate the sampling distribution of coalescence time estimates with a normal distribution for computational convenience. Figure 7 illustrates examples of distributions of estimated coalescence times produced under different mutation rates for a fixed locus and true coalescence time 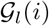, along with the variance approximated under a normal approximation. Further details for the normal approximation are given below.

### 4.2 The composite likelihood

The input for the algorithm are the haplotypes of *M* individuals, from *K* regions in the genome, 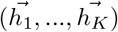, where each 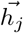 contains *M* haplotypes from locus *j*. We assume that the *K* genes are non-recombining blocks of the genome, and allow free recombination between genes. We allow for each locus to be of different length, and allow for missing characters in the sequences. The rooted gene tree topology, 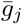, of *M* individuals with branch lengths estimated from haplotypes 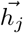 at locus *j* from the pairwise number of differences between the sequences. Each of the *M* individuals must also be assigned to be one of *N* present-day species.

We use a composite likelihood by maximizing the product of likelihoods of each independent gene tree:

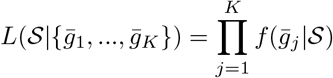

To evaluate the likelihood of an estimated gene tree 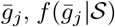, we approximate it by the composite likelihood obtained as products of the individual likelihood functions. For *M* individuals in the tree (*M* ≥ *N*), we decompose the likelihood into *Q* univariate quantities:

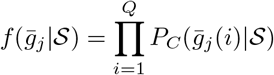

where 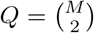 is the number of pairs of individuals in the data set. We index each pair of individuals by a value *i*, (*i* ∈ {1, 2,…, *Q*}), where 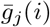 is the estimated coalescence time of pair *i* on gene tree *j*. Note that these *Q* coalescence times are not all independent, as there are only *M* − 1 unique coalescence times on a tree of M individuals.

We model 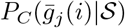 with a zero-inflated MSC-normal hierarchical distribution. Due to the random process of mutation, the frequency of observing zero pairwise mutations at a locus needs to be explicitly modeled, as the MSC-normal distribution does not adequately account for the point mass at zero.

### 4.3 MSC-Normal distribution

For two individuals, *a, b* (indexed by *i*), the divergence time for the species *A, B* respectively (*a* ∈ *A, b* ∈ *B*) is denoted by *τ_AB_*. For a given locus, we estimate a coalescence time 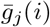 for the pair, based on the estimated local gene tree. We know (assuming no recombination within the locus) that there is some underlying, but unknown, true coalescence time 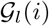.

We model the distribution of location-adjusted true coalescence times, 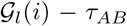, using the coalescent with piecewise constant population size history, with population sizes and times given by 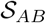. For notation’s sake, we assume the history is a sequence of *R* population size-split time pairs 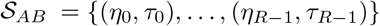, where *η*_0_ = *η_AB_* and *τ*_0_ = *τ_AB_*. At each branch along the tree, we can calculate the likelihood of 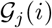 given the coalescence event occurs within the branch 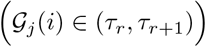, parameterized by the start and end times of the branch, along with the effective population size *η_r_*. To get the overall likelihood of 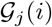, we sum over all the possible branches.

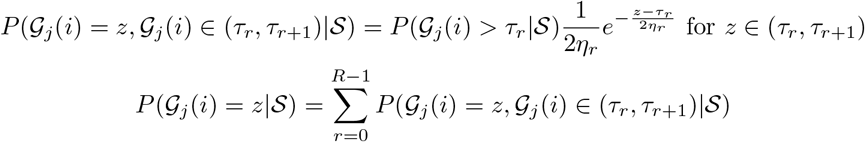

Assuming 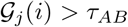, and *τ*_0_ = *τ_AB_*.

Given 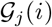, we view the distribution of 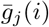 as normally distributed around mean 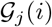, with variance 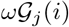, as described earlier.

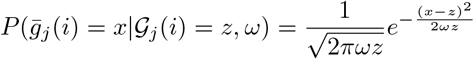

Combining these distributions, we have

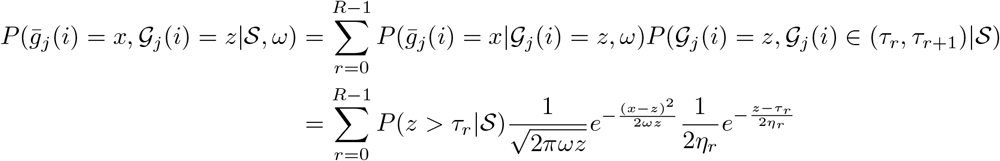

To get the marginal distribution of estimated coalescence times, we need to integrate over the latent variable, 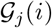, the true coalescence time, which takes values in (*τ_AB_*, ∞)

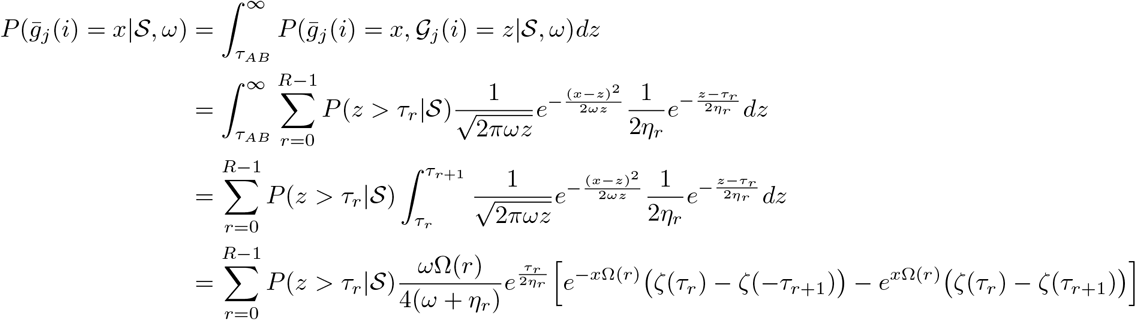

Where

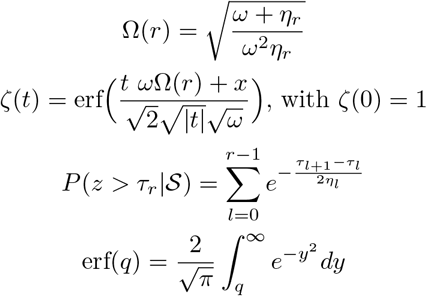

### 4.4 Accounting for no observed mutations

In studying sequence data it is common to encounter genes where two or more individuals have identical sequences, especially when genes are short, or the individuals are of the same species. In constructing a gene tree with no mutations between the two, this pair of individuals would have an estimated coalescence time of 0. For a given pair of individuals (indexed by i on the tree), we can calculate 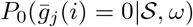, using the MSC and a Poisson distribution of the mutation process. From the Poisson, for a given coalescence time, 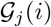, the probability of observing no mutations on the branch of length 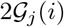 is 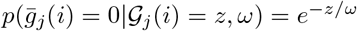.

To obtain the unconditional probability of observing 0 mutations, we need to integrate over all of the possible values of the underlying (and unknown) true gene tree coalescence time, 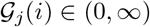:

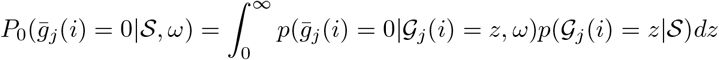

We break the integral into regions of constant population size, indexed by *r* ∈ {0, …,*R* − 1} and evaluate them separately.

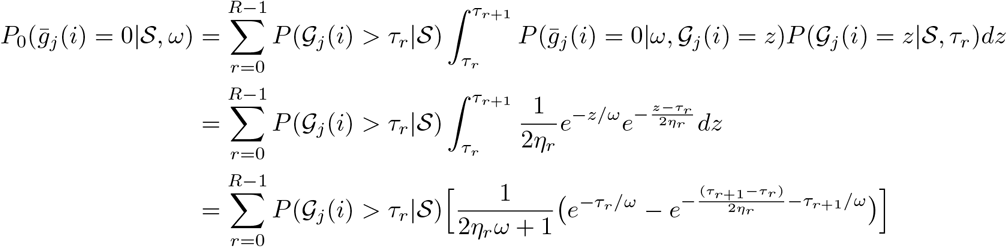

Where *τ*_0_ is the species divergence time for the pair of individuals indexed by i. Calculating the quantity gives us the probability of encountering no mutations between pair i on gene *j* given species tree 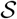, gene length 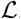, and scaled mutation parameter *θ*. To distinguish this probability from the MSC-Normal distribution also presented above, we subscript the probability with a zero, 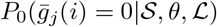, and write the complete likelihood as

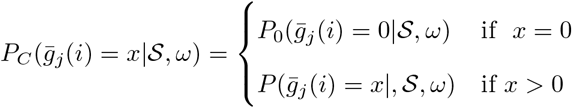

### 4.5 Likelihood weighting

In the composite likelihood, identical information is repeatedly used in multiple probability calculations. For a given node in a gene tree, let *n*_1_ be the number of individuals on one side of the split, and *n*_2_ be the number on the other. Being based on pairwise events, the composite likelihood would then use the information of that node split time *n*_1_ × *n*_2_ times, which can become a large number for nodes deep in a gene tree. We apply a weight to the terms of the likelihood to down-weight this redundant use of information. As we do not observe the gene trees beforehand, we rely on the species tree topology to create the weight values. For a pair of individuals, *i* = (*i*_1_, *i*_2_), *V*(*i*) denotes the split on the tree such that *i*_1_ is on one side of the split, and *i*_2_ is on the other. Given *V*(*i*), denote *n*_1_(*i*) and *n*_2_(*i*) to be the number of individuals on each side of the branch, such that *n*_1_(*i*) × *n*_2_(*i*) is the number of pairs of individuals who share the same split at *V*(*i*). Define weight

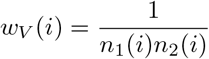

such that, for a given split *V*(*i*),

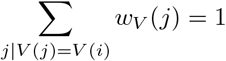

where *j* indicates a pair of individuals (*j*_1_, *j*_2_) that share the same split event *V*(*i*). We apply this weight to each term in the composite likelihood,

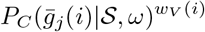

so that the weight of information applied to each split on the species tree is equivalent.

It should be noted that these weights are only used in parameter inference, as using weights which depend on the topology can be problematic when comparing topologies. COAL_PHYRE is able to run with and without the weights applied.

### 4.6 Data simulation

To test the effectiveness of parameter inference of COAL-PHYRE, we conduct simulation studies under varying species tree topologies, divergence times, population sizes, mutation rates, and data set sizes. We simulate gene trees using ms [12] under a bifurcating species tree with piece-wise constant population size and no gene flow or migration after split. For consistency with the assumptions of BPP, we simulate the mutation process using the Jukes and Cantor mutation model [13] through Seq-Gen [26] to produce haplotypes under various mutation rates to introduce varying levels of mutational variance. See Appendix C for more details on the simulations. Although we use a simple model of evolution with a Jukes and Cantor model, performance using other models will likely be similar as long as gene tree estimation is done under the same model as used for simulation.

## 5 Simulation Results

### 5.1 5 species asymmetrical tree

We simulate a tree of 5 taxa, with asymmetric topology (5,(4,(1,(3, 2)))), where species 5 is the outgroup, and 2 individuals sampled per species. The population size within a branch is simulated to be constant, but different between branches, see Appendix section C for exact simulation details. We compare our method, COAL_PHYRE, to BPP [34] and ASTRAL-III [35]. COAL_PHYRE and BPP provide separate estimates of divergence times and population sizes, while ASTRAL-III provides estimates of the coalescence rate of each branch (coalescence rate = branch length/ population size), and does not attempt separate the parameters further. To accommodate the comparatively slow run time of the MCMC-based BPP, we simulate only 100 independent loci for each replicate. It should be noted that COAL_PHYRE can handle much larger sets of genes with only modest increases in run-time. For this data of 5 species, BPP and COAL_PHYRE provide estimates of all 4 split times, as well as the 9 separate population sizes (5 modern-day species and 4 ancestral populations). ASTRAL provides an estimate of 4 external branch lengths, and 2 internal. For each method, we provide as input the known species tree topology, and allow for parameter inference under the true topology. Note that BPP and COAL_PHYRE take as input the sequence data directly, but ASTRAL requires gene trees to be provided. As these simulations use the molecular clock, we use UPGMA (unweighted pair group method with arithmetic mean) to reconstruct gene trees as input to ASTRAL. We simulate under two different mutation rates, *θ* = 0.01, and *θ* = 0.001 (here *θ* = 4*η*_0_*μ* where *μ* is the per generation per base pair mutation rate), representing both high and low levels of mutation, with each locus chosen to be 1000 bp long. Under the *θ* = 0.01 simulation, the the variance in the estimate of coalescence times is higher than for *θ* = 0.01 due to the increased mutational noise.

We simulated 40 separate replicates under the two mutation rates, and used COAL_PHYRE, ASTRAL-III, and BPP to evaluate the accuracy of parameter reconstruction. The results of the estimation from all three methods can be seen in figure 2.

**Figure 2:**
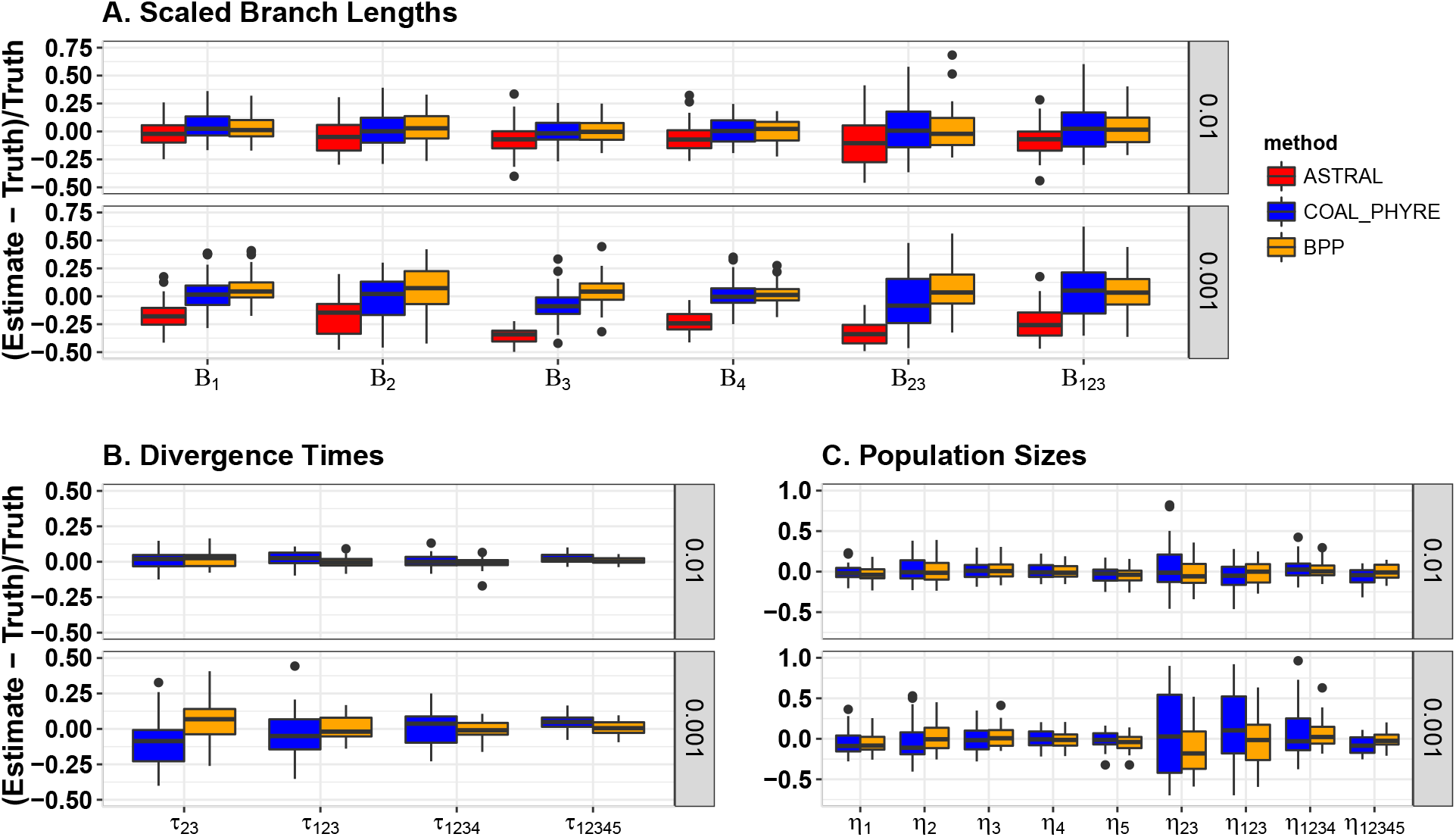
Full parameter estimation for the fixed species tree topology (5,(4,(1,(2,3)))). Comparison of parameter estimates between COAL_PHYRE, ASTRAL and BPP by branch over 40 iterations, using 100 independent loci each iteration. The y-axis gives the standardized deviation from the true parameter value. In each panel, the top plot represents a high mutation rate setting, where mutational variance is low, and the bottom represents a **×** 10 lower mutation rate, where mutational variance is larger. **A)** A comparison of estimated scaled branch lengths (branch length divided by population size) for the three methods. Only branches for which ASTRAL can provide an estimate are included. **B)** A comparison of divergence time estimates between COAL_PHYRE and BPP. **C)** A comparison of population size estimates between COAL_PHYRE and BPP.

We can see that the performance of ASTRAL deteriorates under the low mutation rate model, as the method assumes gene trees are estimated without error, which is violated when the amount of phylogenetic signal in each gene is low. Divergence time estimates are nearly identical between COAL_PHYRE and BPP in the 0.01 mutation rate setting. Under the lower mutation rate, COAL_PHYRE tends to have higher variance and uncertainty in estimating divergence times than BPP. However, it is, similarly to BPP, approximately unbiased. Population size estimates are again nearly identical between COAL_PHYRE and BPP under the 0.01 mutation rate setting. For a lower mutation rate (0.001), the two methods are nearly identical in accuracy for the external population sizes (*η*_1_… *η*_5_) and COAL_PHYRE has more uncertainty than BPP in estimation of internal population sizes, reflecting the well-known challenge of disentangling internal branch lengths from population sizes.

When comparing run times, ASTRAL completed on average in about 1 second per replicate, much faster than either COAL_PHYRE or BPP, but requires pre-computed gene trees before running. COAL_PHYRE outputs results for each replicate in, on average, 1 minute whereas BPP required ~ 10 − 20 minutes to converge, both using a single-core on a standard laptop.

### 5.2 8 species symmetrical tree

Here we simulate a balanced tree topology of 8 species with 2 diploid individuals sampled per species. We simulate under the assumption of constant population size within each branch, but population sizes vary among branches [RN: insert reference to where full details can be found]. Again, we compare COAL_PHYRE to BPP [34], and ASTRAL-III [35]. We simulate 100 independent sequences in each replicate, to compare against BPP at a reasonable run time. Both COAL_PHYRE and BPP can provide estimates of all 7 divergence times, and 15 population sizes (8 modern day, and 7 ancestral). ASTRAL only provides estimates for the leaf population branch lengths, and internal branches which are not directly adjacent to the ancestor of all species in the tree, (so not branch “1234” or “5678”). For BPP and COAL_PHYRE we provide as input the sequence data, the mutation rate, and the known species tree topology. To use ASTRAL, we provide a file of gene trees, pre-estimated using UPGMA, as well as the known species tree topology.

We simulate under two different mutation rates *θ* = 0.01 and *θ* = 0.001 (see above 5 species simulation for discussion on units), with each sequence simulated to be 1000 bp long (Figure 3). Similarly to the 5 species simulation, the branch length estimates of ASTRAL are biased downwards for the low mutation rate setting. As both COAL_PHYRE and BPP explicitly model the mutational noise, they do not experience the same bias. BPP and COAL_PHYRE demonstrate approximately the same level of performance at estimating divergence times and population sizes in the species tree. In particular, both methods provide highly accurate estimates of the leaf branch population sizes (*η*_1_,…*η*_8_). On a single-core laptop computer, COAL_PHYRE completed each of the replicates in 3-10 minutes. We were able to run BPP in approximately 30-60 minutes per replicate. We note that we allow BPP to complete under the recommended settings.

**Figure 3:**
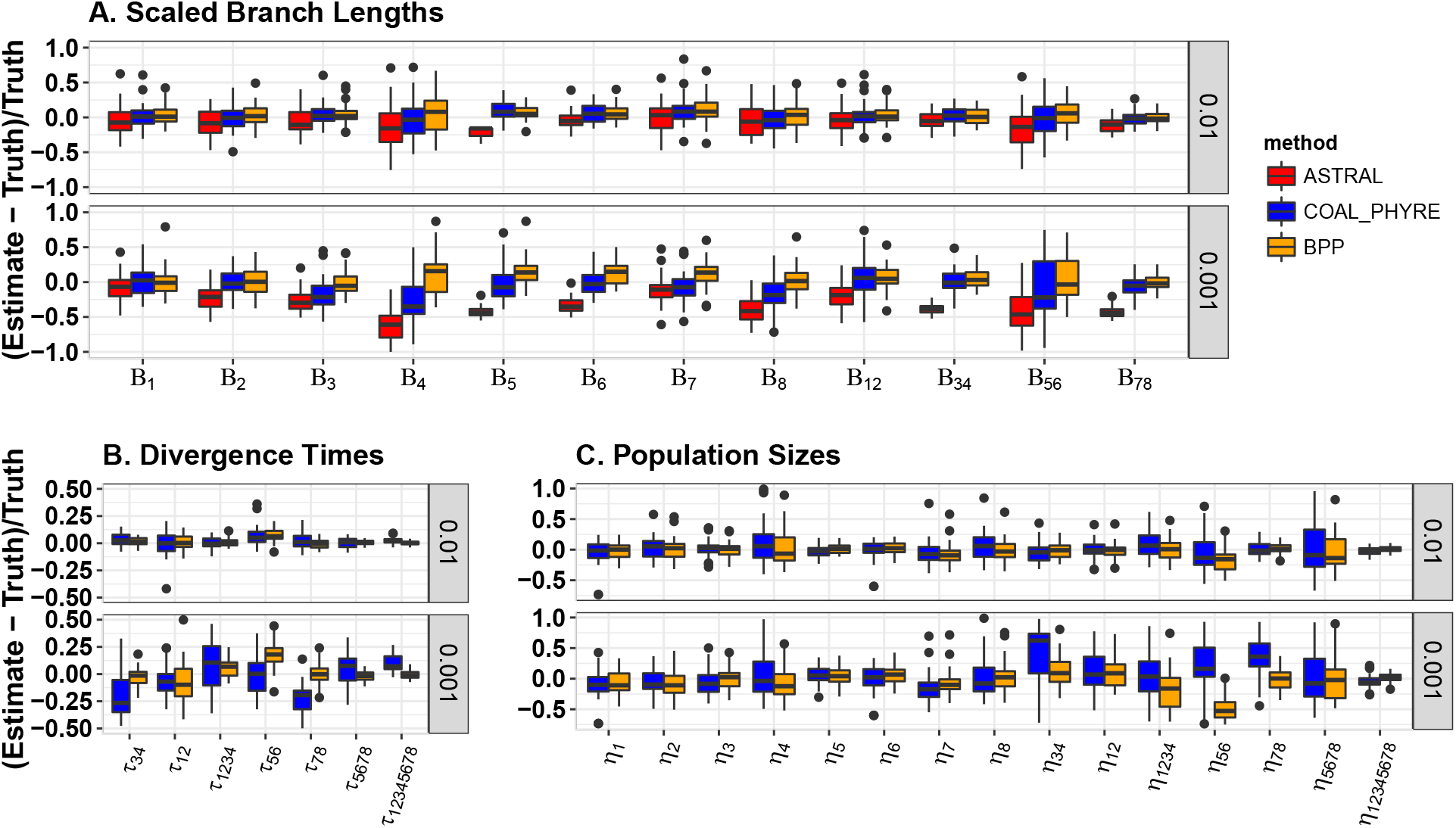
Full parameter estimation for the fixed species tree topology (((1,2),(3,4)),((5,6),(7,8))). Comparison of the parameter estimation accuracy between COAL_PHYRE (blue) and BPP (orange), and ASTRAL-III (red) using 100 independent genes, across 40 independent replicates. **A)** A comparison of estimated scaled branch lengths (branch length divided by population size) for the three methods. Only branches for which ASTRAL can provide an estimate are included. **B)** A comparison of divergence time estimates between COAL_PHYRE and BPP. **C)** A comparison of population size estimates between COAL_PHYRE and BPP.

## 6 Analysis of Gibbon Data

Here we analyze two full-genome data sets from [2] and [33] of four gibbon species: (*Hylobates moloch* (Hm), *Hylobates pileatus* (Hp)), *Nomascus leucogenys* (N), *Symphalangus synd,actylus*(S), and *Hoolock leuconedys*(B). Gibbons (Hylobatidae), close relatives to humans and great apes, are found throughout Southeast Asia’s tropical forests. A recent study, Shi and Yang, 2017[30] (hereby referred to as SY17) used the MCMC program, BPP ([34]), along with a suite of other methods, to attempt to resolve the phylogenetic relationship of these species. The results of the study show there are two most likely species tree topologies, (H, (N, (B, S))), which we will call Tree 1, and (N, (H, (B, S))), denoted by Tree 2. The authors also reported estimates for the population sizes and divergence times on the trees. (Note H= (Hm, Hp) indicating two subpopulations of the *Hylobates* species).

### 6.1 The data

The first data set (Noncoding) consists of 12,413 loci, each of 1,000 bp in length. The second data set (Coding) consists of 11,323 coding loci, each of 200bp in length. Within each data set one human haplotype (O) is used as an outgroup. There are a total of 17 haplotypes at each locus, with two diploid individuals from each Gibbon population, allowing for the estimation of leaf population sizes. See SY17 for a more detailed description of the data.

### 6.2 Results

We use COAL_PHYRE to analyze each of these data sets to provide a likelihood for each of the two topologies, and estimates of the divergence times and population sizes for each tree. To compare with the results of BPP we assume the JC69 [13] model of mutation. As well, we use mutation rate parameters consistent with the means of the Gamma priors used in SY17.

#### 6.2.1 Divergence time and population sizes estimates

The parameter values estimated using COAL_PHYRE, along with those previously estimated in SY17 are presented in Tables 1, 2, 3, 4. In each scenario, we found that COAL_PHYRE assigned the highest likelihood to Tree 1, topology (H, (N, (B, S))), consistent with the findings in SY17. Also, note that population sizes are not reported for the human out group O, as only one haplotype was used, and so no information is available to estimate *η_O_*.

**Table 1:**
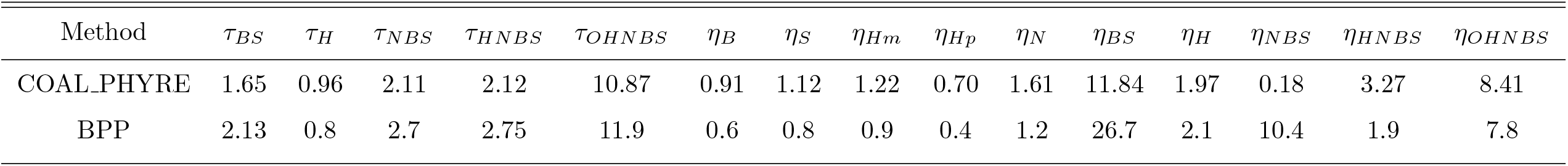
Table of Results: (H, (N, (B, S))) Coding

**Table 2:**
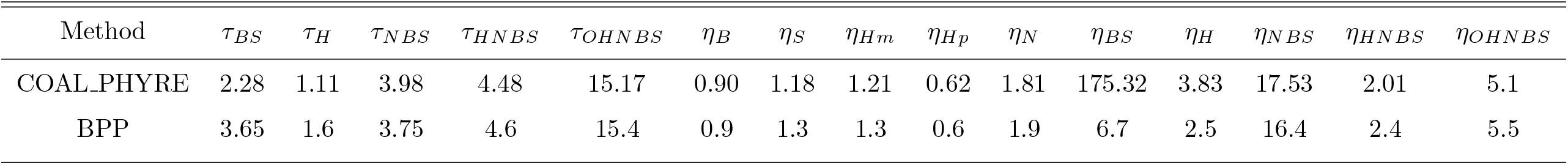
Table of Results: (H, (N, (B, S))) Noncoding

**Table 3:** Table of Results: (N, (H, (B, S))) Coding

| Method | $\tau_{BS}$ | $\tau_H$ | $\tau_{HBS}$ | $\tau_{NHBS}$ | $\tau_{ONHBS}$ | $\eta_B$ | $\eta_S$ | $\eta_{Hm}$ | $\eta_{Hp}$ | $\eta_N$ | $\eta_{BS}$ | $\eta_H$ | $\eta_{HBS}$ | $\eta_{NHBS}$ | $\eta_{ONHBS}$ |
| --- | --- | --- | --- | --- | --- | --- | --- | --- | --- | --- | --- | --- | --- | --- | --- |
| COAL-PHYRE | 1.66 | 1.04 | 1.82 | 2.14 | 10.85 | 0.91 | 1.13 | 1.27 | 0.73 | 1.61 | 3.87 | 1.43 | 24.87 | 3.22 | 8.43 |
| BPP | 1.9 | 1.0 | 3.0 | 3.05 | 11.5 | 0.6 | 0.8 | 0.8 | 0.4 | 1.2 | 22.3 | 2.0 | 2.6 | 1.9 | 7.8 |

**Table 4:** Table of Results: (N, (H, (B, S))) Noncoding

| Method | $\tau_{BS}$ | $\tau_H$ | $\tau_{HBS}$ | $\tau_{NHBS}$ | $\tau_{ONHBS}$ | $\eta_B$ | $\eta_S$ | $\eta_{Hm}$ | $\eta_{Hp}$ | $\eta_N$ | $\eta_{BS}$ | $\eta_H$ | $\eta_{HBS}$ | $\eta_{NHBS}$ | $\eta_{ONHBS}$ |
| --- | --- | --- | --- | --- | --- | --- | --- | --- | --- | --- | --- | --- | --- | --- | --- |
| COAL-PHYRE | 2.29 | 1.12 | 4.16 | 4.45 | 15.17 | 0.9 | 1.18 | 1.22 | 0.62 | 1.82 | 3.73 | 112.32 | inf | 2.00 | 5.10 |
| BPP | 3.75 | 1.6 | 4.3 | 4.8 | 15.25 | 0.9 | 1.3 | 1.3 | 0.6 | 2.0 | 12.5 | 2.3 | 14.4 | 2.5 | 5.5 |

Under the most likely topology (Tree 1) our estimates of the parameters are overall quite similar between coding and noncoding data sets, providing some evidence of internal consistency. To verify this, as suggested in SY17, we fit a regression line, *y* = *bx* between the 5 parameter points (each point a pair of *τ* divergence time estimates, one from the noncoding dataset, the other from coding) to measure the internal consistency of the estimates from COAL_PHYRE. Our analysis under Tree 1 finds *τ*_(*C*)_ = 0.69*τ*_(*NC*)_ with *r*^2^ = 0.988. This demonstrates that our timing estimates are consistent between the two data sets, and that the mutation rate of the coding data is about 2/3 the rate of the non coding loci. SY17 found a rate of 0.73 with *r*^2^ = 0.985, from their analysis. For the population size estimates (*η*’s) of the leaf populations (B, S, N, Hm, Hp) we find *η*_(*C*)_ = 0.95*η*_(*NC*)_ with a correlation of *r*^2^ = 0.995 compared to *r*^2^ = 0.986 from SY17.

We can also compare the correlation between our results and the results from BPP. Divergence time estimates for the (H,(N,(B,S))) coding data set show an *r*^2^ = 0.999 between the divergence times estimated between the two methods, with *τ*_COAL_PHYRE_ = 0.81*τ*_BPP_. For the noncoding data set and tree (H,(N,(B,S)), we find an *r*^2^ = 0.9988 with *τ*_COAL_PHYRE_ = 0.94*τ*_BPP_. When comparing the leaf population sizes we find for the coding data set, *η*_COAL_PHYRE_ = 1.43*η*_BPP_ with *r*^2^ = 0.995. For the noncoding data set we find *η*_COAL_PHYRE_ = 0.97*η*_BPP_ with *r*^2^ = 0.998.

We observe that our parameter estimates overall agree with the results of BPP, differing mainly in estimation of internal population sizes. The largest discrepancies occur on the (N,(H,(B,S)) tree (tree 2), which demonstrates how the two methods handle fitting parameters to a potentially incorrect topology. We acknowledge that SY17 observed BPP had mixing issues for such a large data set, and parameter estimation with short branch lengths can become highly variable. The extremely high population size estimate (which we write as “inf”) of *η_HBS_* in the noncoding tree 2 (N(H(B,S))) indicates that COAL_PHYRE attempts to model extremely high ILS in the HBS branch, attempting to fit a zero-probabilty of coalescence in that interval.

Each of the four tables demonstrates one run of COAL_PHYRE, which on a single core is able to run on average in 10(±5) hours. As reported in SY17, BPP took approximately 200 hrs for each analysis on a single core using the same data as COAL_PHYRE.

#### 6.2.2 Predicted distribution of estimated coalescence times

Parameters on the species trees are estimated to best match the distribution of estimated coalescence times in the data, according to some likelihood function. In this section we assess the fit of the predicted distribution of estimated pairwise coalescence times of the Gibbon data when using the zero inflated MSC-Normal distribution implemented in COAL_PHYRE.

For a given set of tree parameters (topology, times and population sizes), we can study the resulting marginal distributions of estimated times. As we have two sets of tree parameters for each scenario, one from each method, we can compare the distributions predicted by each against the distribution of estimated times from data.

We specifically study the most likely tree topology, Tree 1 (H,(N,(B,S))), parameterized by the sets of divergence times and population sizes from Tables 1 and 2 (see Figures 5 and 4, respectively). Using the parameter values estimated by both methods, we can compare the predicted distribution under each set of parameters against the actual sampled distribution from the estimates across loci, and against one another to assess a level of ‘best fit’ to the data.

**Figure 4:**
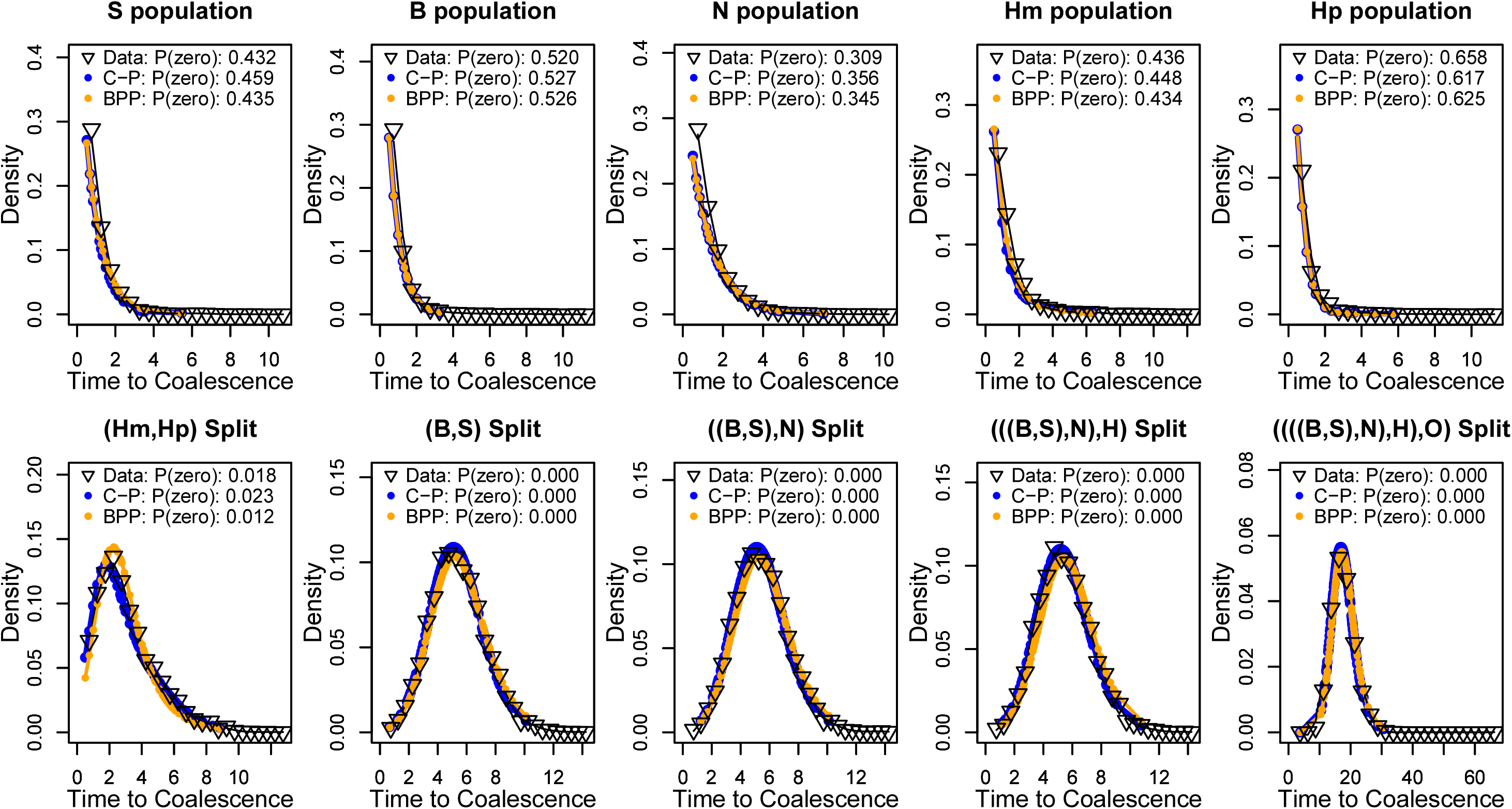
Distribution of Estimated Coalescent Time: Tree 1 of Noncoding Gibbon data. Comparison of zero-inflated MSC Normal distributions of estimated coalescence times using parameters inferred by COAL_PHYRE (Blue) and BPP (Orange) along with the distribution of estimated coalescence times from the data (black triangles) for each proposed split event. The top row: for two individuals sampled from the same population (as indicated by the plot header), the distribution of the estimated coalescence times. The bottom row: for a given split event, e.g. “((B,S),N) Split”, the distribution plots the estimated time to coalescence for an individual sampled from the (B,S) subset, and one from the (N) subset. The figure legend also includes the observed (or predicted) fraction of zeros in the data.

**Figure 5:**
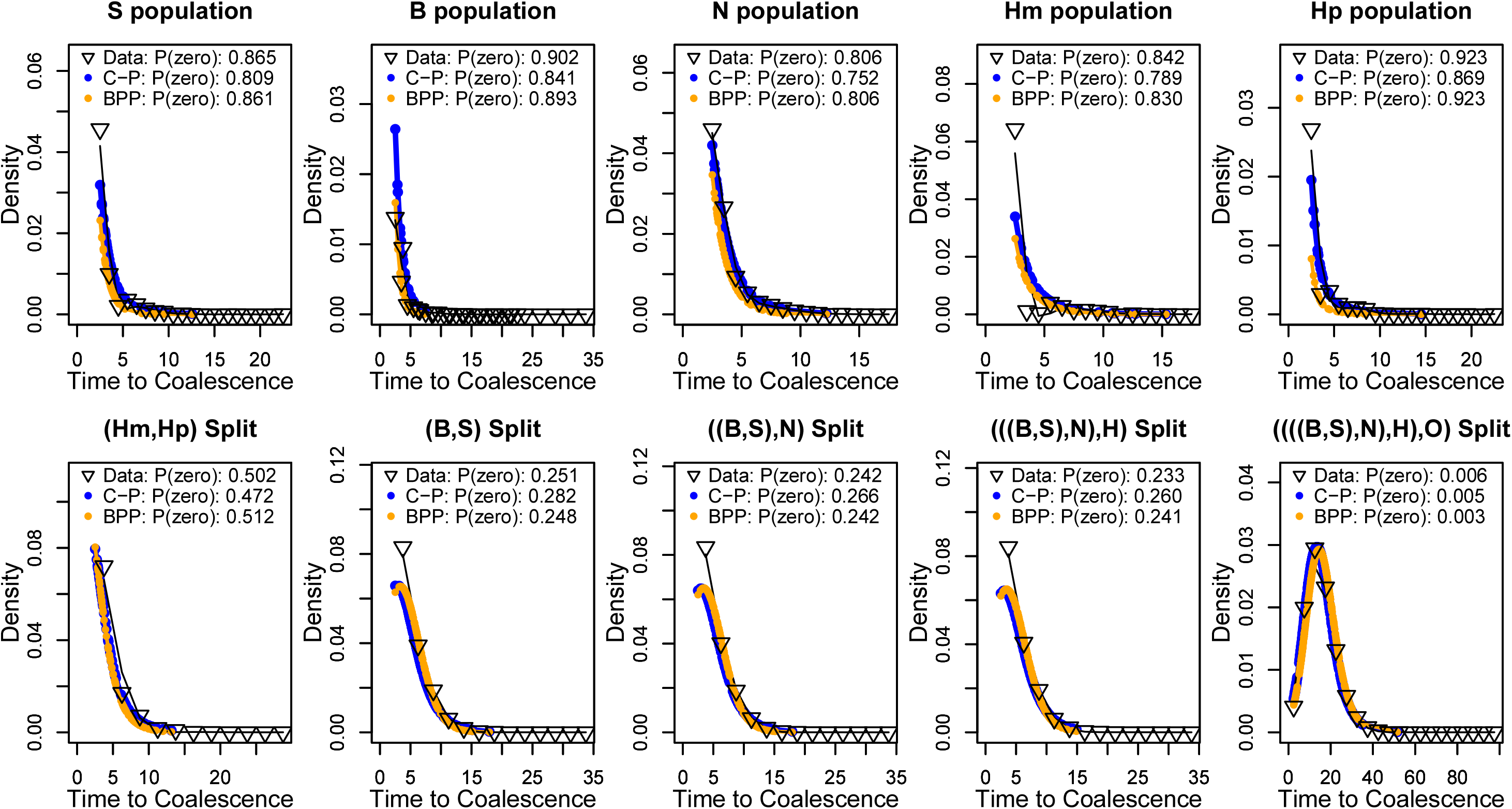
Distribution of Estimated Coalescent Time: Tree 1 of Coding Gibbon data. Comparison of zero-inflated MSC Normal distributions of estimated coalescence times using parameters inferred by COAL_PHYRE (Blue) and BPP (Orange) along with the distribution of estimated coalescence times from the data (black triangles) for each proposed split event. The top row: for two individuals sampled from the same population (as indicated by the plot header), the distribution of the estimated coalescence times. The bottom row: for a given split event, e.g. “((B,S),N) Split”, the distribution plots the estimated time to coalescence for an individual sampled from the (B,S) subset, and one from the (N) subset. The figure legend also includes the observed (or predicted) fraction of zeros in the data.

Figure 4 shows the distribution of binned estimated pairwise coalescence times from the data, along with the predicted distributions using the parameters of both COAL_PHYRE and BPP for the noncoding data set under Tree 1. From the plot, we can see that the predicted distributions between the two methods agree almost exactly in each panel. Figure 5 is the same approach, using the coding dataset.

Across all distributions of estimated coalescence times, it is expected that COAL_PHYRE should fit the data as well or better than the parameters from BPP, as the parameters inferred by COAL_PHYRE are estimated to fit specifically this likelihood.

Each plot also shows the predicted fraction of sequences that have no pairwise differences, as well as the observed frequency of zeros in the data. Comparing the parameters from COAL_PHYRE and BPP on the accuracy of predicting the fraction of zeros shows that BPP is slightly more accurate in this respect, on average.

Overall, the parameters inferred by each method fit the shape of the distribution of estimates well.

#### 6.2.3 Run times

Each of the four tables demonstrates one run of COAL_PHYRE, which on a single core is able to run on average in 10(±5) hours. As reported in SY17, BPP took approximately 200 hrs for each analysis on a single core using the same data as COAL_PHYRE.

## 7 Discussion

Our simulations show that COAL_PHYRE provides estimates that are comparable to BPP and much more accurate than estimates obtained using ASTRAL-III. We observe a strong effect of mutational variance on estimates obtained using ASTRAL in a low mutation rate setting. An advantage of COAL_PHYRE is that it is straightforward to separate the effects of the two genetic processes that generate the input data by studying the role of both the MSC and the normal distribution.

For the Gibbon data set, we showed that our method can analyze genomic-sized data sets with similar performance to BPP, with an order of magnitude decrease in run time. The composite likelihood approach of only using pairwise coalescence times implemented and presented here seems to sufficiently capture the relevant parts of the data needed to infer the tree parameters. COAL_PHYRE recovered the same most likely topology as presented in [30], for both the coding and non coding datasets. The largest discrepancies between our method and BPP in the analysis of the gibbon data was in fitting parameters to tree 2, which both methods infer to be an incorrect topology. We also see that large deviations in parameter estimates can have negligible effect on the estimated distribution of estimated coalescence times, for example *η_BS_* in Table 2, and the resulting effect in Figure 4.

When studying species tree estimation, it is typical to also study topology reconstruction accuracy. We have found in our simulations that ASTRAL is superior in topology reconstruction, and with the speed of ASTRAL compared to COAL_PHYRE, we do not make claims that our method is superior for inferring topologies. The information extracted and used from the data by the two methods is largely orthogonal; ASTRAL uses purely the topological information from each estimated gene tree, and discards all information on coalescence times, whereas COAL_PHYRE only uses marginal coalescence times from each gene, and discards topology information. This lends itself to the idea that the information used in COAL_PHYRE and ASTRAL can be combined or that, at least, be employed in tandem. We also acknowledge work done in [1, 24] which presents a data pre-processing step to counter the effects of mutational variance for programs such as ASTRAL which do not directly model it.

Lastly, none of these methods account for migration/gene flow between species after divergence, something which is common in most real data sets. Failing to account for this potential gene flow can affect topology inference as well as drastically effect divergence time and population size estimation. Accounting for and modeling potential sources of admixture is a next step for these parameter inference methods. It is worth noting that a preprint for an extension of BPP implementing the full MSC with introgression (MSci) has recently been released [7]. Identifying locations of admixture and fitting admixture branches to a species tree are left to future work for COAL_PHYRE.

More studies are needed to understand the robustness of the different methods, for example with regards to substitution models or, and in particular, the effect of recombination within a block. Genomic data is not truly composed of free recombining segments with no internal recombination, which is effectively assumed by all methods analysed in this paper. To address the problem of recombination within blocks, a potential approach is to divide blocks into even smaller units, thereby increasing the amount of mutational variance within each unit, but decreasing the probability of recombination within the unit. As COAL_PHYRE is designed specifically to handle increased variance in estimation, this could be a potential work-around in cases where recombination might be a challenge.

## 7.1 Software Availability

Along with this manuscript, we provide code (implemented in C++) available for download which implements the likelihood presented here, named COAL-PHYRE. The code is implemented in C++ and freely available at https://github.com/gaguerra/COAL_PHYRE.

## Appendices

## A Notation Reference

- *N*: Number of species considered.
- 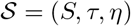: A species tree parameterized by topology *S*, split times *τ*, and population sizes *η*.
- *K*: Number of independent loci/genes.
- *M*: Number of sampled individuals (*M* ≥ *N*).
- 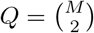 : Number of pairs of individuals.
- 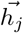: Set of *M* haplotypes at locus *j*, *j* ∈ {1, 2,…, *K*}.
- 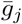: Estimated gene tree at locus *j*.
- 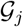: True gene tree at locus *j*.
- *a, b*: Individuals sampled from populations *A, B*, respectively.
- *τ_A,B_* : The split time of species *A* and *B* according to 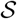.
- Each pair of individuals are indexed by an integer *i*,in (1,…, *Q*).
- 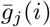: The estimated coalescence time of pair *i* at locus *j*.
- 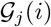: The true time to coalescence of pair *i* at locus *j*.
- *μ*: The per generation per base pair mutation rate.
- 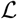: The number of base pairs of gene.
- θ: The population scaled mutation rate, *θ* = 2*μη*_0_, for the reference population size, *η*_0_
- 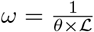
- 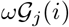: Mutational estimation variance of the true coalescence time.

## B Fit of the MSC-Normal distribution

In this paper we have discussed that estimated coalescence times can be modelled with two sources of variance, one from the coalescent process, and the other from mutational process. In figure 6 we visualize an example of the approximation to the distribution of estimated coalescence times obtained using the MSC-Normal distribution. As well, we plot the true coalescence times when times are known without error. From the figure we notice that in the presence of estimation error, estimated coalescence times can be more recent than the species divergence time. The MSC-Normal distribution acts as an approximation to the distribution of estimated coalescence times, and captures this tail of recent times, as well as the overall distribution.

**Figure 6:**
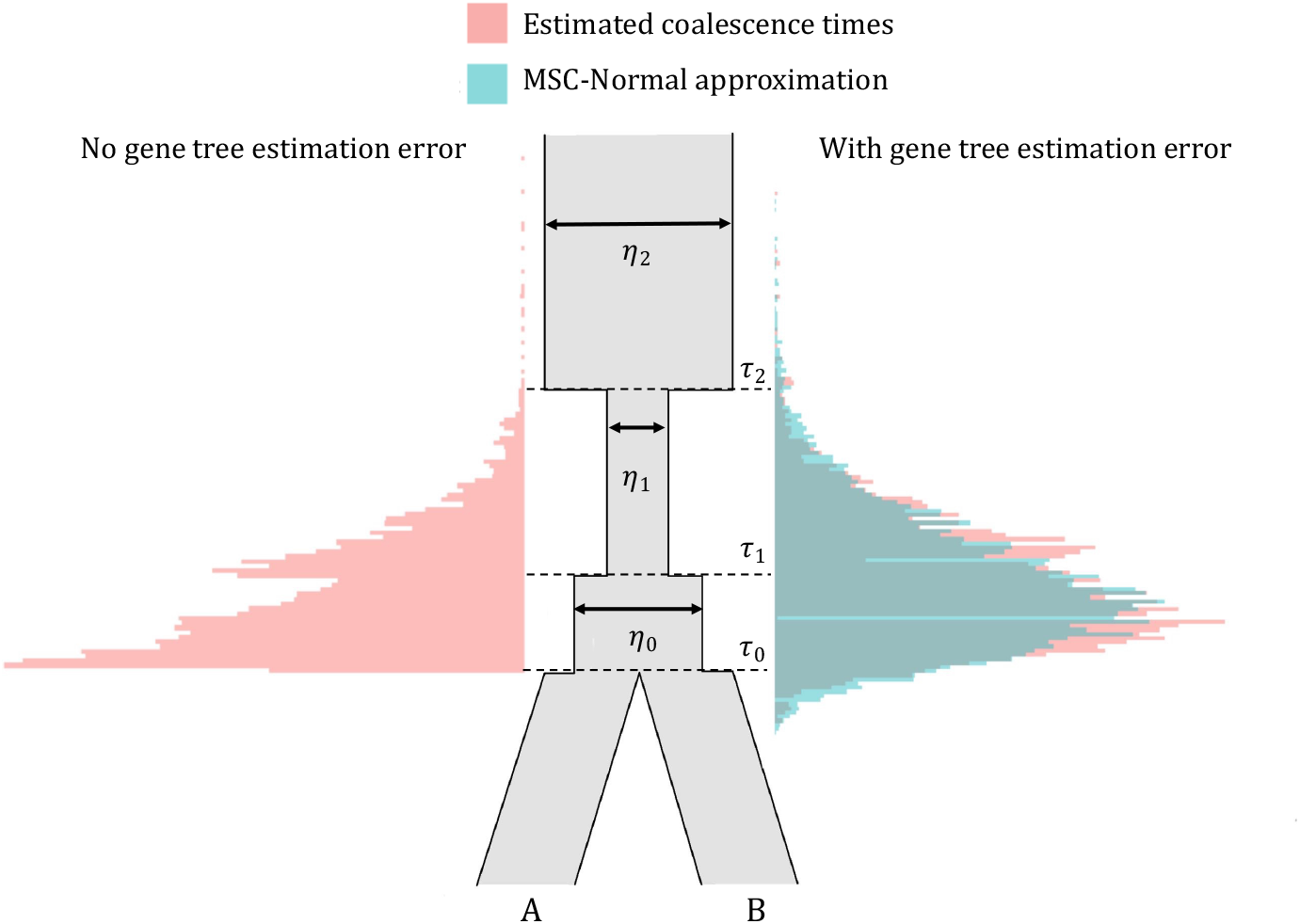
MSC-Normal approximation of coalescence times with coalescence estimation error. Center: Species tree of two individuals with a piece-wise constant population size history. On the left: Plotted in red is the distribution of coalescence times of these two individuals under simulation when the true gene trees are known, so no estimation variance is present. On the right: Plotted in red is the distribution of estimated coalescence times of the same two individuals from simulated sequence data, where variance is introduced due to the mutational process. In green is the MSC-normal approximation to the distribution of estimated coalescence times using the known tree parameters (τ’s and η’s) and mutation rate *θ*.

## C Further Simulation Details

### C.1 5-species simulation details

In this simulation study we analyzed a species tree of 5 species (labeled 1…5) with 10 individuals (labeled 1…10) where 2 individuals are from each species (i.e individuals 1 and 2 are from species 1). We simulate the rooted species topology (5,(4,(1,(2,3)))).

For a single replicate, we use ms to generate K independent gene trees of 10 individuals, 2 from each species, and Seq-Gen [26] to generate sequence data from the gene trees. To generate K = 100 gene trees of 10 individuals with species labeled as integers 1 through 5:

~~~
./ms 10 100 -T -I 5 2 2 2 2 2 -n 1 1.8 -n 2 2.4 -n 3 1.0 -n 4 2.0 -n 5 3.0 -ej 1.0 2 3
-en 1.0 3 2.4 -ej 1.5 1 3 -en 1.5 3 3.0 -ej 2.2 3 4 -en 2.2 4 4.0 -ej 4.0 4 5
-en 4.0 5 5.0 | tail +4 |grep -v // >gene.trees
~~~

In ms [12], time is measured in units of 4*η*_0_ generations, whereas COAL_PHYRE measures time in 2*η*_0_ generations, so that times from COAL_PHYRE must be halved to compare to the units of ms. As well, population sizes in ms are diploid, whereas in COAL_PHYRE we measure population sizes as haploid. To compare with ms, population sizes from COAL_PHYRE need to be doubled.

From the **gene.trees** file, and for a given mutation parameter *θ* (which we used either 0.01 or 0.001 in our simulation), and sequence length 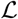, we use Seq-Gen. For example, for *θ* = 0.001 and 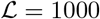:

~~~
./Seq-Gen -mHKY -l 1000 -s 0.001 <gene.trees >seqfile
~~~

We use this **seqfile** file as input to COAL_PHYRE.

### C.2 8-species simulation details

For 8 species, 2 individuals sampled per species, we generated a single replicate of K = 100 independent gene trees using:

~~~
./ms 16 100 -T 8 2 2 2 2 2 2 2 2 -n 1 1.5 -n 2 2.5 -n 3 2.0 -n 4 6.0 -n 5 0.5 -n 6 1.0 -n 7 3.0 -n 8 4.0 -ej 0.5 2 1 -en 0.5 1 6.0 -ej 0.75 4 3 -en 0.75 3 1.0 -ej 0.8 8 7 -en 0.8 7 2.0 -ej 1.3 6 5 -en 1.3 5 4.0 -ej 1.5 3 1 -en 1.5 1 5.0 -ej 1.8 7 5 -en 1.8 1.5 -ej 2.0 5 1 -en 2.0 1 6.0 | tail +4 | grep -v // >gene.trees
~~~

For *θ* = 0.01 and gene length 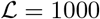, we generate sequence data with Seq-Gen:

~~~
./Seq-Gen -mHKY -l 1000 -s 0.01 <gene.trees >seqfile
~~~

We use this **seqfile** file as the input to COAL_PHYRE.

## D Normal Approximation To Poisson, A Simulation

Throughout we discuss the distribution of estimated coalescence times. The estimation error from the mutation process, conditional on a branch length, follows a Poisson distribution. As our estimated coalescence times are not discrete, we use the Normal approximation to the Poisson. In this section we demonstrate in a simple simulation scenario, that this approximation is well fit to model the estimation error. For a given coalescence time (fixed here to be 5 in units of 2*η*_0_ generations), we simulate 1000 pairs of sequences, of length 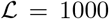 base pairs, under varying scaled mutation rates, *θ* (indicated in figure 7 legend), to generate an empirical distribution of estimated coalescence times from the number of pairwise differences. Figure 7 shows these distributions versus the normal approximation presented earlier. This demonstrates the accuracy and suitability of the Normal approximation to mutational variance in time estimation.

**Figure 7:**
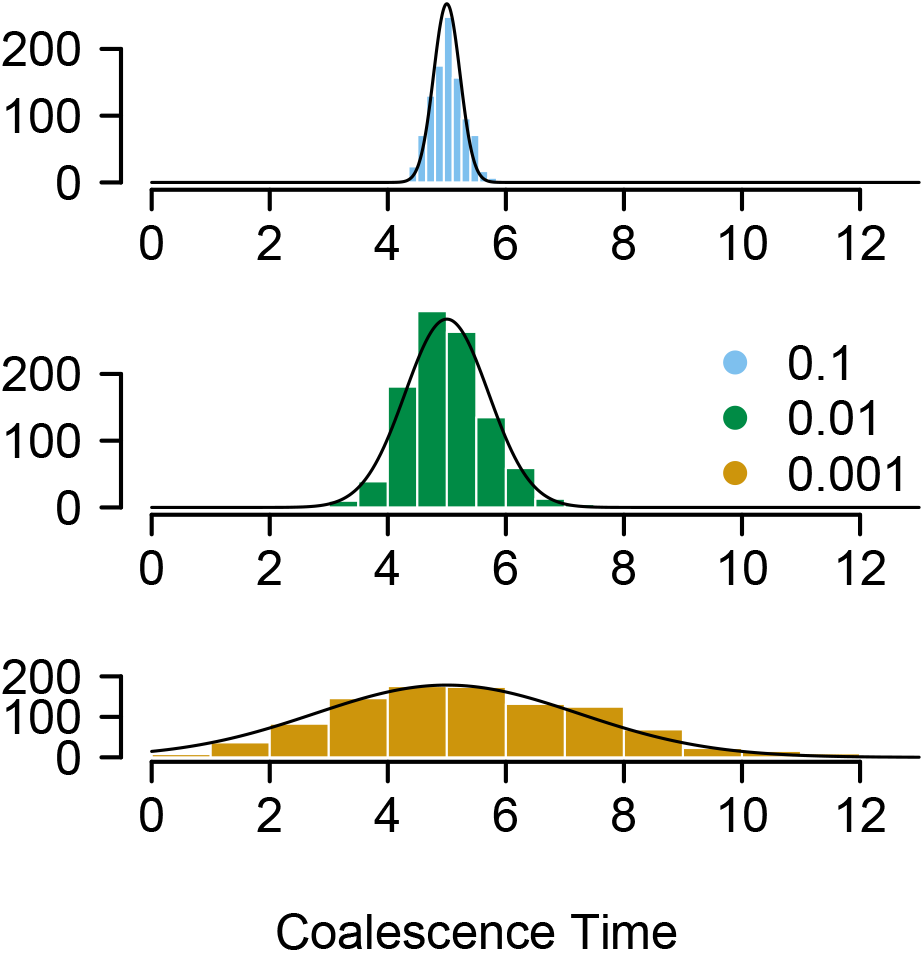
Modeling Mutational Variance: The Normal approximation to Possion variance in coalescent time estimation error due to the mutational process.

## Notes

### Competing Interest Statement

The authors have declared no competing interest.

